# Force Propagation in Active Cytoskeletal Networks

**DOI:** 10.1101/2025.08.13.669984

**Authors:** Shichen Liu, Rosalind Wenshan Pan, Heun Jin Lee, Shahriar Shadkhoo, Fan Yang, David Larios, Chunhe Li, Hao Wang, Zijie Qu, Rob Phillips, Matt Thomson

## Abstract

Conventional materials science establishes clear structure–function relationships: grain boundaries modulate strength by blocking dislocations, and defect density limits conductivity through electron scattering. Active matter presents a fundamentally different challenge, components continuously consume energy to generate forces, creating a dual identity as both structural elements and force-generating machines. This material–machine duality raises a fundamental question: which structural parameters govern functional behavior in systems where constituents actively generate forces rather than merely responding to external inputs? Here we demonstrate that percolation is the critical mechanism that transforms active cytoskeletal networks from energy-dissipating materials into work-performing machines. Using light-controlled microtubule-kinesin networks, we show that increasing bundle length from 0.9 µm to 5 µm triggers a percolation transition that enables global force organization. Networks below the threshold remain incapable of performing coordinated work, while networks above it develop correlation lengths exceeding 240 µm, generate 25-fold stronger forces, and extract 1000-fold more mechanical work, powering transport of live cells across 800 µm distances. Network simulations reveal that introducing just 5-10% longer bundles creates a connected component containing 90% of microtubules, establishing the precise threshold that drives this material-to-machine transition. Our findings establish percolation as the fundamental mechanism governing whether active matter systems function as passive materials or as coordinated machines—providing design principles for both synthetic and biological systems. increasing bundle length from 0.9 ***µ***m to 5 ***µ***m increasing bundle length from 0.9 µm to 5 µm

## Introduction

From metals to metamaterials, engineers treat microstructure as a dial that quantitatively sets bulk response: shrinking aluminium grains from millimetres to micrometres multiplies yield stress[1, 2], ten-nanometer superlattices slash silicon’s lattice thermal conductivity[3, 4], and re-entrant honeycombs drive Poisson’s ratio below −0.8[5–8]. In every case a geometric or compositional feature becomes a reliable design rule. Active materials upend that rulebook. Because every filament, motor, or microbe continuously burns fuel and injects forces, the network remodels within seconds to minutes, and static gauges such as mesh size or cross-link density—so powerful for equilibrium solids—often fail to predict how an active gel will behave—limiting these systems to scientific curiosities rather than deployable materials[9–14].

To harness energy-consuming active matter we must pinpoint the threshold at which random stresses reorganise into coordinated mechanical work. Which structural parameter can predict that crossover? *Past work shows that traditional descriptors—mesh size, filament density, packing fraction—fail to predict when energy input reorganises an active network into coherent motion*. Reconstituted actin networks with identical mesh size behave as soft elastic solids at low myosin occupancy but liquefy when motor activity rises, even though filament length and bulk density vary little [15]. This solid-to-fluid transition coincides with motor forcing (*≈*0.1–1 s) overtaking the *≥*10 s viscoelastic relaxation of the cross-linked mesh [16]. Mitotic spindles stiffen only after antiparallel microtubule overlaps percolate along the *≈*50 *µ*m body [17–19], and confined *Bacillus subtilis* suspensions reorganise into single vortices while packing fraction stays near 0.4 [20]. No accepted metric—robust to continuous turnover—links structure to the onset of coherent work across these examples.

Using a reconstituted microtubule kinesin system, we show that bundle length controls whether active microtubule networks perform useful work or dissipate energy locally. Short bundles (0.83 *µ*m) create disconnected patches with velocity correlations extending 57.4 *µ*m, while longer bundles (4.7 *µ*m) form connected networks that organize forces over 247.5 *µ*m. Simulations show that when 10% of microtubule bundles exceed 5 *µ*m, the network exhibits a percolation transition where 90% of the network becomes connected. The transition to connected networks amplifies cargo-transport forces 25-fold from 1 pN to 26 pN and increases extractable work by three orders of magnitude. The transition enables optical control of cargo transport, allowing light-directed movement of cells and particles over millimeter distances. We establish a structure-function relationship for active matter—demonstrating that bundle length drives functional transitions of active networks from force dissipating to force propagating through percolation.

## Results

### Active matter incubation time switches contraction between the global force-propagating phase and the local force-dissipating phase

During the cycles of growth, reconfiguration, and division, cells dynamically switch their internal cytoskeletal networks between functioning as passive structural materials and active force-generating machines through precise modulation of the length and bundling effect of microtubules [13, 21–24]. To investigate the transition between passive and active properties of the two cytoskeletal organizations, we used a defined and optically controlled in vitro experimental system composed of stabilized microtubules and engineered kinesin motor proteins [25–29]. Our microtubule kinesin active matter system consists of stabilized microtubules labeled with fluorophore and kinesin motor proteins fused with optically dimerizable iLID (improved light-induced dimer) proteins [30]. Upon light activation, the kinesin protein dimerization induces interactions between kinesins and neighboring microtubules, which leads to microtubule network formation and contraction into asters. In previous work, we observed that pre-incubation of motors and microtubules could induce a transition between long-range (one aster spanning the entire light-illuminated region) and short-range phases (many disconnected asters in the light-illuminated region) of organization. We hypothesize that this transition likely stems from changes in microtubule bundling, a key factor in determining network properties and sizes [27, 28]. Bundles can significantly alter the effective length and rigidity of microtubule structures [9], potentially serving as a mechanism for switching between organizational phases. Our in-vitro system allows us to isolate and study how bundle formation influences the critical transition between passive material states where forces remain localized and dissipated, and active machine states capable of propagating forces globally to perform mechanical work. Therefore, we performed a systematic study of system organization while quantitatively varying the duration of motor-microtubule incubation time.

We investigated how pre-activation incubation time affects cytoskeletal network organization in a light-controlled active matter system. Our experiments revealed two distinct phases of microtubule organization and dynamics. For incubation times below 40 minutes, we observed a local phase characterized by small microtubule asters smaller than 50 *µ*m confined within the light-activated region. In contrast, incubation times exceeding 65 minutes led to a global phase with long-range effects, forming structures up to 800 *µ*m and recruiting microtubules from over 300 *µ*m away from the light activated region [31]. We established a calibration curve by calculating the average intensities at different microtubule densities based on tubulin concentration in our system. Quantitative analysis showed that the global phase outperforms the local phase in material concentration and transport. The global phase exhibited an increase in microtubule density of 3.4-fold within the light-activated region, compared to a small increase of 1.3-fold in the local phase (Figure 1B and Supplemental Figure 3). Moreover, the global phase induced large-scale microtubule network buckling and long-range material recruitment, capabilities absent in the local phase. Our findings demonstrate that extended incubation enables the emergence of coordinated, system-wide behaviors in active microtubule networks, significantly enhancing their ability to concentrate and organize material over large distances.

**Fig. 1.**
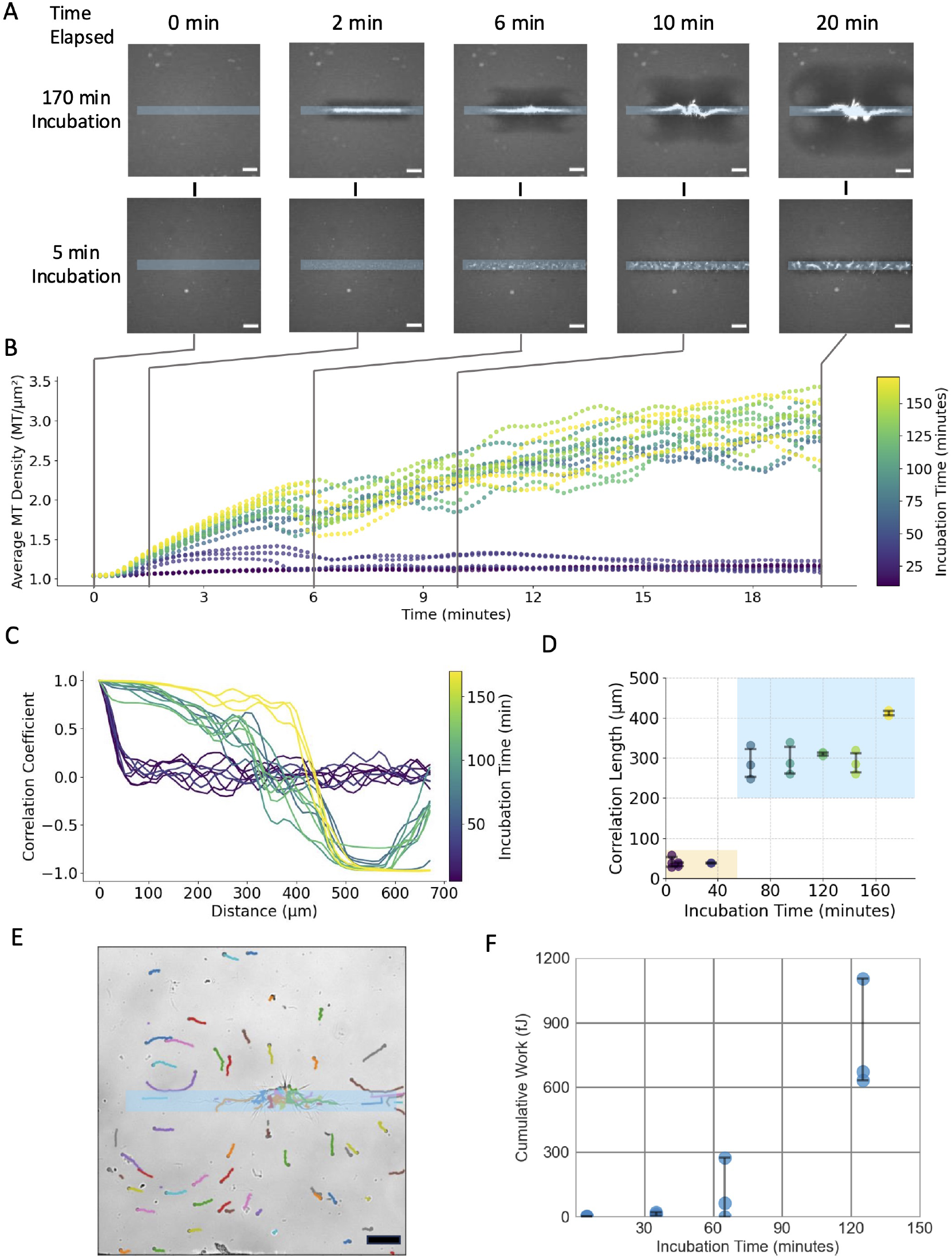
Microtubule Incubation-Induced Phase Transition Between Local and Global Phases. **A**, Images of labeled microtubules during aster assembly. The top row shows the global contraction phase with an incubation time of 170 minutes, while the bottom row shows the local contraction phase with an incubation time of 5 minutes. Scale bar: 100 *µ*m. **B**, Microtubule density in the light-illuminated region, color-coded by incubation time. We observed increases in microtubule density in the light illuminated region as time progresses in the experiments with higher incubation time, while microtubule density stayed at 1-1.3 for experiments with lower incubation time. **C**, Euclidean two-point correlation coefficient of the light-illuminated region, color-coded by incubation time. We computed the correlation coefficient by fixing a point at the right most point of the light illuminated region and varied the second point at different distances. Experiments with incubation time over 65 minutes showed persistent correlated movement within the light activated region, and a negative correlation representing directional change of velocity changes. While the experiments with incubation time below 65 minutes showed a sharp drop-off at 50 *µ*m showing no persistent correlated movements. **D**, Correlation length computed from the correlation coefficient function. The orange shaded region represents the local contraction phase, and the blue region represents the global contraction phase. Correlation length *l* of the local phase is calculated by fitting *A·*exp(*−x/l*) to the correlation coefficients. Correlation length *l* of the global phase is calculated by finding the distance between two points when the correlation coefficients decayed by 1*/e* **E**, Representative particle movement in the global force-propagating phase. Scale bar: 100 *µ*m. **F**, Active matter system performs more than 1,000 times more extractable mechanical work after incubation by moving polystyrene tracer beads. Work calculated by *W* = 6*πηrvd*, where *v* is measured using particle tracking. N = 15 beads for 5 minutes incubation, N = 17 beads for 35 minutes incubation, N = 18 beads for 65 minutes incubation, N = 14 beads for 125 minutes incubation, n = 3 independent experiments for each incubation time.

To measure how far the coherent movements extend through the system, we analyzed spatial correlations in the velocity field. We first computed the velocity field form particle image velocimetry (PIV) of the aster contraction images. Then, we calculated the Eulerian two-point correlation coefficient (Figure 1C) and the corresponding correlation length (Figure 1D) [32]. This spatial correlation coefficient reveals the persistence of velocity vector directions and magnitudes across space. We computed the correlation coefficient by anchoring the first point at the rightmost edge of the rectangular light pattern and varying the second point’s distance horizontally along the light-activated region (Figure 1A). For the local phase, we fitted *A·*exp(−*x/l*) to the correlation coefficient data, where *l* represents the correlation length. For the global phase, due to its complex behavior and negative correlation, we defined the correlation length as the characteristic length scale at which the correlation coefficient decayed by 1*/e*.

Our analysis revealed two distinct correlation lengths for the global and local phases. The local phase, observed in experiments with incubation times under 40 minutes, exhibited correlation lengths clustering below 57.4 *µ*m (Figure 1D). This aligns with our observations of small asters with uncorrelated motions. We observed an abrupt increase in correlation length from 57.4 *µ*m to 247.5 *µ*m at 65-minute incubation time, marking the shift from local to global phase. Within the global phase, correlation lengths increased gradually from 247.5 *µ*m to 411.3 *µ*m with longer incubation times. Hence, we classify the local phase as having correlation length below 80 *µ*m and the global phase as having correlation length above 230 *µ*m (Figure 1D). We did not observe any further discontinuous jumps once the system entered the global phase. The local phase exhibits a sharp decline in the correlation coefficient as inter-point distance increases, eventually fluctuating around zero at 60 *µ*m (Figure 1C). In the global phase, we observed persistently correlated motion over long distances, with the correlation coefficient becoming negative at 300-500 *µ*m (Figure 1C), approximately half the length of the microtubule network. This transition to negative correlation indicates opposing velocity vectors, reflecting symmetric material recruitment.

The microtubule network transitions from a passive material unable to transport cargo to an active machine capable of directed transport precisely at the boundary between local and global correlation phases, as demonstrated by tracking 10 *µ*m polystyrene tracer beads in solutions activated with identical light patterns across different incubation times. In samples incubated for less than 40 minutes (local phase, correlation length *<* 57.4 *µ*m), beads exhibited only Brownian motion and remained effectively stationary relative to their starting positions, confirming that forces generated by motors dissipated locally without producing coordinated mechanical output. In stark contrast, samples incubated for more than 65 minutes (global phase, correlation length *>* 247.5 *µ*m) actively transported beads across distances up to 200 *µ*m from their initial positions (Figure 1E). We estimated a minimum force of 26 pN acting on the beads (a 25-fold increase) by analyzing their displacement trajectories using Stokes’ law, corresponding to an increase in extractable mechanical work from nearly zero to 1103 pJ (Figure 1F). This calculation provides a conservative lower bound as it accounts only for fluid drag resistance and not additional forces required to reorganize the microtubule network structure. The directed transport demonstrates that the percolation transition we identified enables the system to propagate forces across macroscopic distances, transforming from a passive material that dissipates energy to an active machine capable of performing extractable mechanical work. The capacity to transport cargo emerges only when the network forms a connected giant component spanning hundreds of micrometers, providing direct experimental evidence for our proposed material-machine duality in active cytoskeletal networks.

### Microtubule bundle size drives a percolation transition that enables global force propagation

To elucidate the microscopic mechanism underlying the transition from the local force-dissipating phase to the global force-propagating phase, we performed fluorescence recovery after photobleaching (FRAP) experiments. FRAP allows us to extract an effective bundle length — i.e. a mobility-based proxy rather than a direct length measurement — from the diffusion coefficient of fluorescent microtubules as they repopulate a bleached region. We found that the length of the microtubule structures increased linearly from 0.9 *µ*m to 5 *µ*m over a period of 2 hours, with a rate of 2.69 *µ*m/hr at room temperature (Figure 2A). This rate of increase is 4 times higher than the GMP-cpp (Supplemental Information) polymerization rate of tubulin, which is 0.5 *µ*m/hr at room temperature [33]. GMP-cpp is a slowly hydrolyzable analog of GTP that stabilizes microtubules, and its polymerization rate serves as a reference for the observed growth rate. Thus, the higher growth rate observed in our experiments suggests that microtubules exhibit bundling behavior, which increases the effective length of each microtubule structure by 5-fold in 120 minutes.

**Fig. 2.**
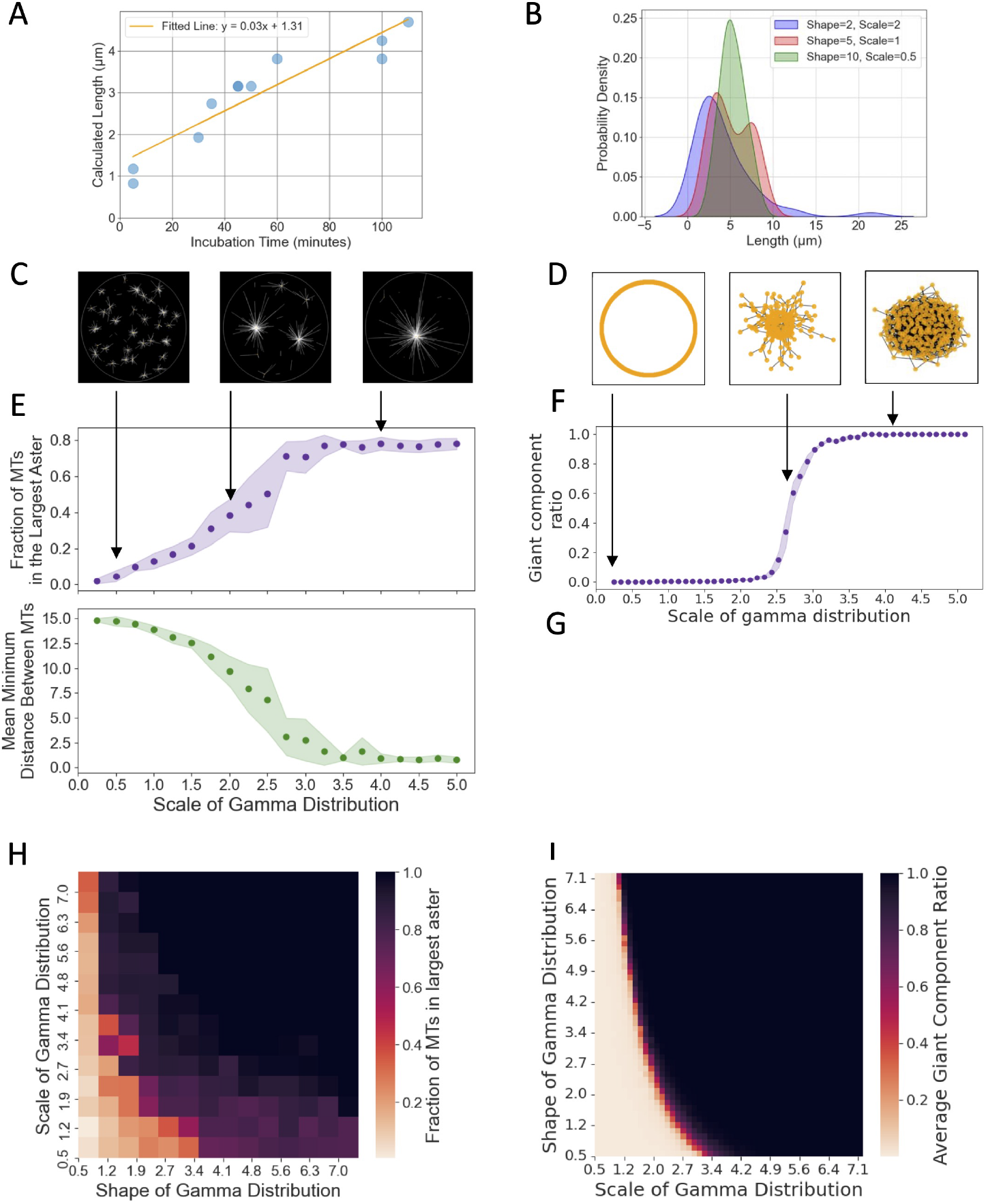
A fivefold increase in microtubule bundle length enables force propagation phase in active networks.. **A**, Incubation time increases microtubule effective length at the rate of 2.69 *µ*m per hour measured by Fluorescence Recovery After Photobleaching (FRAP) **B**, Kernel density estimates (KDE) of microtubule length distributions drawn from a modified Gamma distribution, where each segment length is constrained to 1-25 *µ*m and the total length does not exceed 200 *µ*m. Three parameter sets (shape=2, scale=2), (shape=5, scale=1), and (shape=10, scale=0.5) illustrate how these constraints alter the probability density profiles relative to a standard Gamma distribution**C**, Images of representative Cytosim simulations under different microtubule length distribution. **D**, Images of representative percolation simulation under different microtubule length distribution. **E**, The top plot shows the fraction of microtubules in the largest aster versus the scale parameter of the gamma distribution. The bottom plot shows the mean of the minimum distance between each pair of microtubules versus the scale parameter of the gamma distribution. Shape = 1 **F**, Giant component ratio vs scale of gamma distribution for the percolation simulations. Shape = 1 **G**, Representative gamma distribution for simulated microtubule length. **H**, Heatmap showing the relationship of the shape and scale of gamma distributed microtubule species and fraction of microtubules in the largest aster. **I**, Heatmap showing the relationship of the shape and scale of gamma distributed edge species and the giant component ratio.

To validate that these changes in microtubule length affect system organization, we first employed Cytosim, a numerical simulation platform designed to model cytoskeletal mechanics by simulating each microtubule and kinesin (Figure 2C) [34]. We investigated whether increasing the length of the microtubules would cause the system to transition from having many local asters to having a global aster. The length of the microtubules was modeled using a modified gamma distribution, where the shortest microtubule was 1 *µ*m, and the longest was 25 *µ*m [35] (Figure 2B). We fixed the total length of microtubules in simulations to 1000 *µ*m, so as microtubules grow longer, the total number of microtubules decreases. As we increased the scale parameter, which introduced a small portion of long microtubules, the simulated active matter system transitioned from having many isolated local asters to having a global aster. We quantify connectivity with the percolation order parameter *S* = *N*_GC_*/N*_tot_, the probability that a randomly chosen filament belongs to the giant component. When the scale parameter of the microtubule length distribution increased to 2.75, with an average microtubule length of 3.7 *µ*m, we observed plateaus in both the order parameter and the mean distance between any pair of microtubules, indicating the system’s transition into the global phase (Figure 2E).

To generalize beyond particle detail, we built a graph model in which nodes represent filament centers of mass and edge probabilities scale with filament length, rooted in percolation theory [36] (Figure 2D). This abstraction ignores filament orientation and steric exclusion, offering a mean-field baseline that captures connectivity statistics without micromechanical detail. Because the edge probability depends only on bundle length and not on spatial distance, the system is analytically equivalent to an Erdős–Rényi (ER) random graph once those probabilities are fixed. As we increased the scale parameter to introduce longer microtubules, we observed a sudden, dramatic increase in network connectivity: the size of the largest connected component jumped until all nodes merged into a single “giant component” spanning the system (Figure 2F). We quantify this jump with the same percolation order parameter we used for qunatifying the cytosim simulations *S* = *N*_GC_*/N*_tot_; in the simulation *S* climbs from 0.1 to 0.9 scale parameter of the microtubule length distribution increased to 2.75 (Figure 2F). Mean-field finite-size scaling predicts *S ∝ n*^*−β/ν*^ *f* [(*c −* 1)*n*^1*/ν*^] with *β* = 1 and *ν* = 1*/*2 (Supplementary Eq. 3). Plotting *S n*^1*/*3^ versus (*c −* 1)*n*^1*/*3^ (with *c* the mean degree) for graph sizes from 6.4 k to 102 k nodes collapses the data onto the Erdős–Rényi mean-field curve, yielding critical exponents *β ≃ γ ≃* 1 (Supplementary Fig. 7)[37]. This giant component represents a continuous path of connectivity through which forces propagate across the entire network.

Scanning both the scale and shape parameters of the length distribution confirms this mean-field percolation boundary: our simulated system flips sharply from a disconnected network with many small isolated clusters to a connected network dominated by a single giant component (Fig. 2H). Introducing even a small fraction of extended microtubules is sufficient to cross the threshold. Below it, the network remains fragmented into islands where forces dissipate locally; above it, the spanning cluster creates a continuous path hundreds of micrometers long through which forces propagate globally. This connectivity-driven transition directly enables the directed transport and mechanical-work extraction observed in our experiments, providing a quantitative explanation for how subtle changes in microtubule bundle size trigger dramatic shifts between passive-material and active-machine states.

### Controlled aster movement demonstrates improved efficiency of the active matter machine

To study the long-range interactions enabled by the global force-propagating phase of the active cytoskeleton network, we constructed an aster merger operation (Figure 3A) where we connected two asters using light patterns, as previously described by [30]. In both the local and global phases, two asters were formed by circular light-excitation patterns, with the local phase aster being smaller and depleting less material compared to the global phase aster. Upon connecting the pair of asters using a rectangular pattern with a high aspect ratio, we observed that in the global phase, the two asters merged after a short delay time (1 minute), while the aster pair in the local phase did not exhibit any movement. These findings further reinforced that the moving and merging global phase asters are dynamic and constantly remodeling, whereas the local phase asters are closer to steady-phase structures [38].

**Fig. 3.**
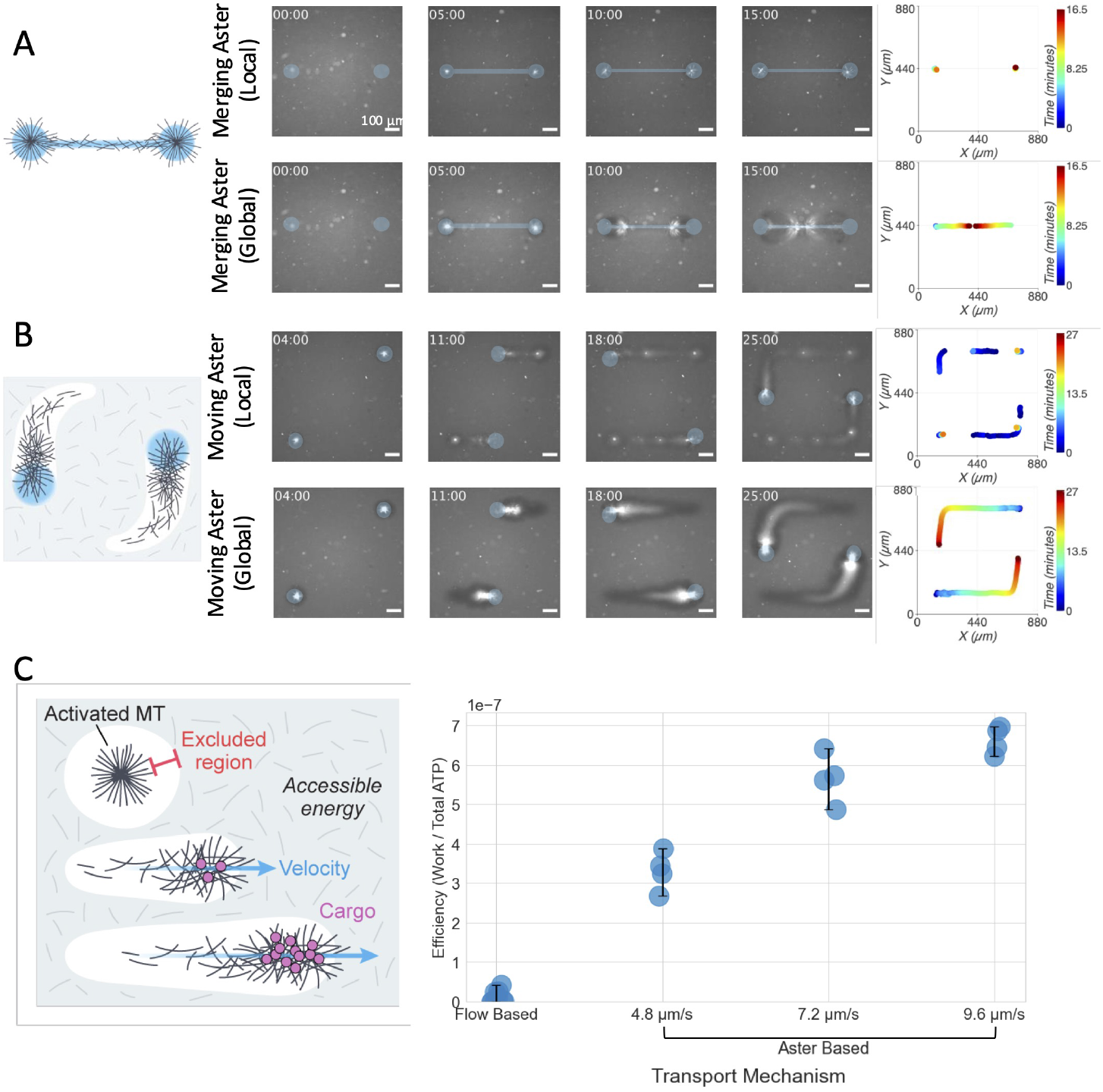
Force propagation driven by global contraction enables transport of microtubules and enhances work efficiency. **A**, Aster linking in both local (top) and global (bottom) phases. Plots on the right shows trajectories of asters position using image thresholding. The asters in the local phase do not exhibit movement when linking them with an additional thin light pattern, while the two asters in the global phase merged. **B**, Aster moving in both local (top) and global (bottom) phases. The plots on the right shows the time each aster persists. Asters in the local phase did not follow the light pattern, thus many new asters emerged as the dynamic light pattern moved. Asters in the global phase followed the light pattern throughout the entire movement of the dynamic light pattern. **C**, Left: Schematic illustration of our experimental approach for measuring work efficiency, showing activated microtubules, the excluded region for ATP calculation, aster movement velocity, and cargo transport. Right: Quantification of transport efficiency (work output/ATP consumption) comparing flow-based transport with aster-based transport at increasing velocities (4.8, 7.2, and 9.6 *µ*m/s). Aster-based transport shows up to 14-fo1ld1 higher efficiency that increases with movement speed. Time unit: minutes

The ability to direct aster formation and movement, both spatially and over time, marks a critical progression toward transport applications using the active cytoskeleton network. We demonstrated the simultaneous movement of multiple asters using dynamic light patterns (Figure 3B). As asters moved, inflows of microtubule bundles emerged within the light pattern, feeding into and pulling the aster behind. Concurrently, outflows, resembling comet-tail streams, trailed the moving asters. During the aster movement, the global phase asters followed the dynamic light patterns without interruption. In contrast, local phase asters struggled to match the pace of the light patterns, often leading to transient and static asters that subsequently gave rise to new aster formations along the light pattern’s path.

We leveraged this aster-based transport mechanism to significantly enhance work extraction efficiency in our active cytoskeletal system. By first creating an aster that captures tracer beads and then moving the light pattern, we could transport both the beads already incorporated in the aster and recruit additional beads along the path. This aster-based approach improved the efficiency of the system by 14-fold compared to non-aster-based (flow-based) transport, with efficiency further increasing as aster movement speed increased from 4.8 *µ*m/s to 9.6 *µ*m/s (Figure 3C). We calculated efficiency as the ratio of mechanical work performed (estimated via Stokes’ law) to ATP consumed in the microtubule-depleted region. Similar to our earlier force calculation, this calculation represents a conservative lower bound because: (1) Stokes’ law only accounts for fluid drag resistance and not the additional forces required for network reorganization, thus underestimating the total work performed; (2) a significant portion of ATP is consumed for internal network maintenance and reorganization rather than directly contributing to bead transport; and (3) we do not account for the additional work performed in transporting the microtubule material itself along with the beads. Despite these conservative estimates, the dramatic efficiency improvement demonstrates how the percolation transition fundamentally transforms the network from a passive material to an active machine with enhanced force propagation and work extraction capabilities that scale with the system’s operating parameters.

### Force propagating active matter powers cell transporter and droplet motility

The active network’s ability to generate force and transport materials suggests its potential as an engine and transport agent for biological processes and future bio-robots. To demonstrate this capability with biological materials, we investigated the movement of suspended human Jurkat cells within the active cytoskeleton network.Upon light activation, a contracting global phase aster formed and began capturing cells. Using dynamic light patterns (Figure 4A), we directed the aster to move 1 mm while collecting cells along its path (Figure 4B). As the aster moved, its cross-sectional area increased linearly (Figure 4C), aiding in cell retention. When approaching a cell, the aster’s inflows generated power in the range of 10 to 59 nW, driving the cell toward its center (Figure 4D). The global phase aster demonstrated the ability to capture multiple cells simultaneously. Initially, it only captured cells within the light-activated region. However, as incubation time and aster size increased, material outside this region was rapidly recruited, expanding the aster’s capture range. Of 32 cells captured, only 2 were released during transport, both from the aster’s “arms”. Cells in the aster’s core were consistently transported throughout the experiment, suggesting structural differences between the “arms” and “core”. The core’s randomly crosslinked structure may provide greater carrying capacity, due to the more rigid scaffold to keep cells inside the aster core, compared to the arms. Notably, cells that were not captured showed minimal movement compared to captured cells (Figure 4E). The precise control and localized force generation capabilities of our active matter system, operating in the 10-59 nW range, making it well-suited for delicate tasks such as single-cell manipulation, enabling the gentle transport or positioning of individual cells within microfluidic devices without the risk of photodamage or electrical disruption from other methods [39, 40].

**Fig. 4.**
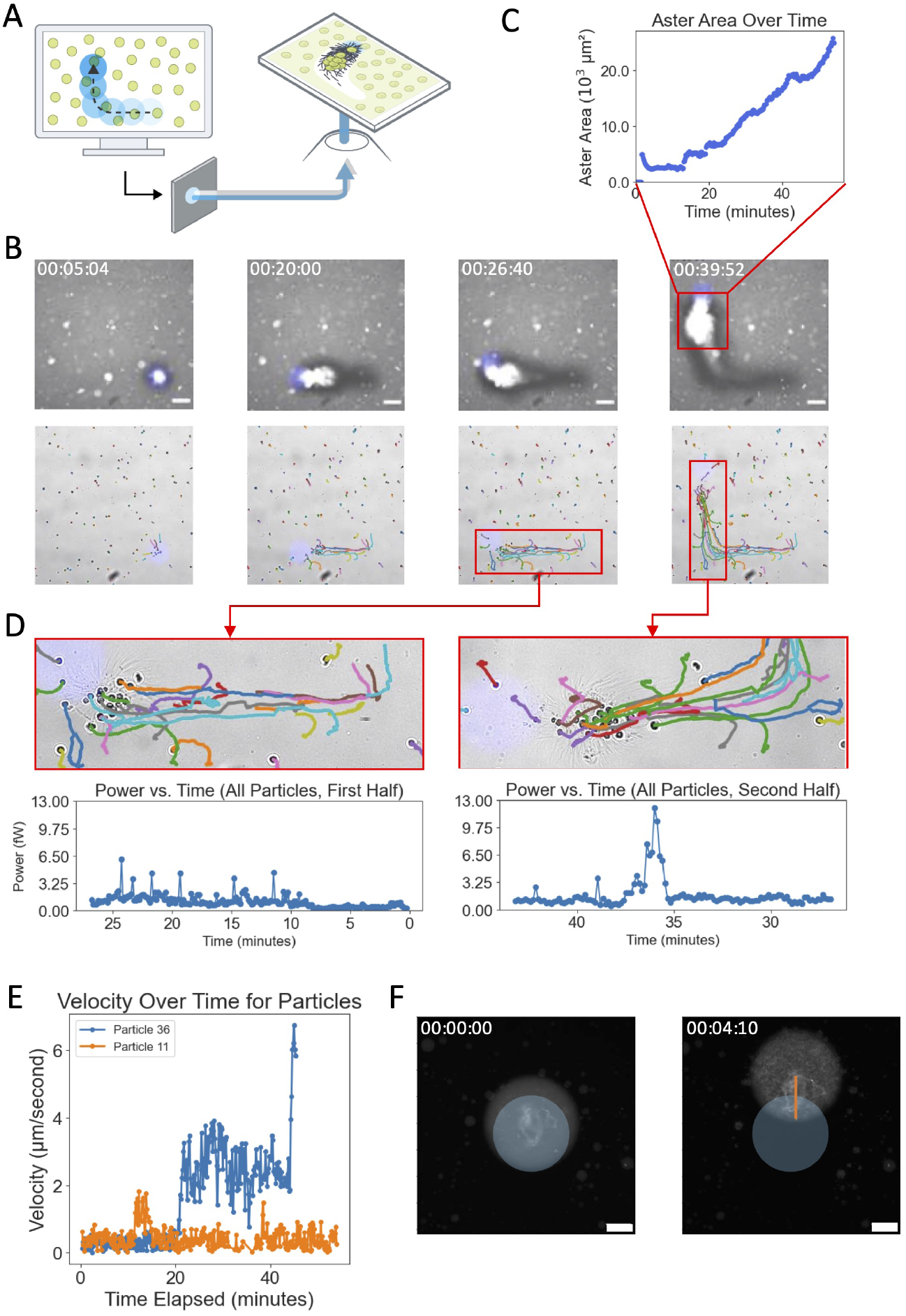
Aster-based active cell transport. **A**, Schematics of cell transport use active matter. Briefly, in a field with active matter, and cells, we utilized a dynamic light pattern similar to the one in Figure 3B to recruit and transport the cells using global phase asters **B**, Time-lapsed images of using aster (top) to concentrate and transport Jurkat T cells (bottom). Scale bar, 100*µ*m.**C**, Aster area increases as aster displacement increases due to constant recruitment of microtubules. **D**, Power generated by moving cells with respect to time. Briefly, we calculated the force using the same method as in Figure 2B by using the Stokes equation and cell tracking. Power is calculated by *P* = *W/*Δ*t* **E**, Example of cell velocity of a stationary cell (orange) vs a cell captured by the aster (blue). **F**, Proof of concept motile droplet powered by contracting active cytoskeleton. Blue light indicates the light illuminated region and orange line represents the trajectory of the droplet. Scale bar, 100 *µ*m

In the biological context, the cytoskeleton and the active fluids generated by it have been proposed to lead to the emergence of cytoplasmic streaming, which can enhance cellular transport. To mimic this biological process, we encapsulated our active cytoskeleton network in aqueous droplets emulsified in oil. When activated by light, the active cytoskeleton network did not seem to propel the aqueous droplet in either the global phase or the local phase (Figure 4F). However, outside of our current defined phases, we were able to generate directed movement of the droplet by light activation, and the movement quickly stopped when we turned off the light. In this phase, the active network does not contract into global or local asters but exhibits a highly crosslinked state with minimal contraction. We hypothesize that the crosslinked network created localized tension at the water-oil interface, which resulted in Marangoni flow and propelled the droplet forward.

## Summary and Outlook

Our results demonstrate how a modest increase in microtubule bundle length drives a mean-field percolation transition, identifiable by a sudden jump in the giant-component order parameter *S* > 0.9. After the percolation transition, velocity correlations stretch four-fold, and bead-tracked forces rise 25-fold, turning the network from a local energy sink into a micrometer-to-millimeter–scale actuator. FRAP shows that the growth rate of free microtubules exceeds pure GMP–CPP polymerization, implicating motor-mediated bundling. Cytosim simulations and a Γ-threshold graph model reproduce the same order-parameter jump and collapse onto Erdős–Rényi scaling (*β* ≈ *γ* ≈ 1). Because long bundles randomize connectivity, the critical length grows only logarithmically with network size: a ten-fold larger device would need just 10–20 % longer bundles.

Our percolation analysis treats bonds as effectively permanent on the motor-forcing timescale, but far richer physics—and new control opportunities—should emerge once motor forcing and bond lifetime become comparable: connectivity thresholds may shift, re-enter, or even cross over into directed-percolation universality. Exploring this *active-percolation* regime is compelling because it marries deep theory to tangible design. First, it tests how nonequilibrium noise renormalizes classical exponents or spawns new fixed points. Second, it couples geometry to dynamics, turning the giant component into a living scaffold whose lifetime statistics can be tuned. Third—and most important for applications—it provides a blueprint for materials that stiffen, relax, or self-heal on demand: by adjusting bond lifetimes with light, ATP, or engineered cross-linkers, one could dial not just *when* a network gels but *how long* it stays gelled, programming cycles of load bearing and fluidization. Systematically mapping the order parameter *S* across bundle length and bond lifetime would therefore elevate active percolation from a theoretical curiosity to a practical framework for building adaptive, reconfigurable active-matter devices.

Cells may exploit the same sparse-connector knob: modest shifts in filament length or cross-link kinetics could let spindles, lamellipodia, and tissue-scale actomyosin sheets toggle between compliant exploration and forceful contraction as mechanical demands change. Similar connectivity effects have been reported in other driven networks—flagellar bundles in dense bacterial suspensions, field-polarised Janus colloids, and active emulsions—which develop system-spanning clusters once a minority of long-range connectors persist longer than they break. In all these cases, activity remains local until connectivity percolates; thereafter, energy can be channelled into coordinated, system-wide work. Recognising this mechanism therefore offers a unifying lens for living assemblies and emerging synthetic active materials.

In summary, our work shows that adding a sparse tail of long filaments is the decisive lever that flips an active network from a passive energy sink into a machine-like actuator, and the critical length grows only logarithmically with system size. This gives engineers a calculable—rather than empirical— recipe for the minimal structural tweak required at any footprint. The same connectivity principle can, in principle, be adopted by cytoskeletal hydrogels, active-nematic films, field-polarised colloids, bacterial collectives, and living cell–laden scaffolds. Near-term applications range from soft-robotic laminates that stiffen or relax on demand, to microfluidic conveyors and pumps that redraw their own flow paths with light[41, 42], to adaptive wound dressings that alternately bear load and fluidize for tissue regeneration. Scientifically, the next frontier is to chart the full landscape where structure meets activity—varying filament length, bond lifetime, and dimensionality—testing whether the mean-field exponents we observe persist, become re-entrant, or cross over to directed-percolation regimes under dynamic bond control, and then pushing the concept into fully three-dimensional fabrics and living tissues. Mastering this connectivity-driven switch therefore opens a route to the first generation of *living metamaterials*—systems that integrate sensing, actuation, and self-repair in a single, continuously powered network.

