## Supplemental information for "Force Propagation in Active Cytoskeletal Networks"

August 13, 2025

### 1 PIV and Optical Flow Methods

#### 1.1 Optical Flow

Optical flow is a method used to estimate the motion of objects between consecutive frames of a video sequence by analyzing the apparent motion of brightness patterns in the images. The Farneback method is a specific optical flow algorithm that uses polynomial expansions to estimate the velocity field.

Code implementation in optical_flow.ipynb

##### 1.1.1 Optical Flow Assumptions

The basic assumption of optical flow is the brightness constancy constraint, which states that the intensity of a pixel remains constant between consecutive frames. Mathematically, this can be expressed as:

$$I_1(\mathbf{x}) = I_2(\mathbf{x} + \mathbf{d}(\mathbf{x})), \tag{1}$$

where $I_1$ and $I_2$ are the intensities of the pixel at position $\mathbf{x}$ in the two frames, and $\mathbf{d}(\mathbf{x})$ is the velocity vector of the pixel.

##### 1.1.2 Farneback Method

The Farneback method approximates the velocity field using polynomial expansions. It represents the local neighborhood of each pixel by a polynomial, which can be expressed as:

$$I(\mathbf{x}) \approx \mathbf{a}_0 + \mathbf{a}_1 x + \mathbf{a}_2 y + \mathbf{a}_3 x^2 + \mathbf{a}_4 xy + \mathbf{a}_5 y^2, \tag{2}$$

where $\mathbf{a}_i$ are the coefficients of the polynomial.

To estimate the velocity field, the method constructs an over-determined system of linear equations by sampling the polynomial coefficients in a neighborhood around each pixel. This system is then solved using least squares minimization to find the velocity vector $\mathbf{d}(\mathbf{x})$ that best fits the brightness constancy constraint.

##### 1.1.3 Velocity Field Calculation

Once the velocity field $\mathbf{d}(\mathbf{x})$ is obtained, the velocity field $\mathbf{v}(\mathbf{x})$ is calculated by dividing the displacement by the time interval $\Delta t$ between frames:

$$\mathbf{v}(\mathbf{x}) = \frac{\mathbf{d}(\mathbf{x})}{\Delta t}. \tag{3}$$

#### 1.2 Particle Image Velocimetry (PIV)

Particle Image Velocimetry (PIV) is a technique for measuring fluid flow velocity fields by tracking the displacement of tracer particles between sequential image frames.

##### 1.2.1 PIV Processing Parameters

In our analysis, PIV was conducted using a three-pass algorithm to enhance the accuracy and resolution of the velocity field. Each pass refines the previous estimate by using smaller interrogation windows and step sizes, effectively increasing spatial resolution. The parameters used for each pass are:

- First pass: Interrogation window of 64 pixels, step size of 32 pixels.
- Second pass: Interrogation window of 32 pixels, step size of 16 pixels.
- Third pass: Interrogation window of 16 pixels, step size of 8 pixels.

These parameters are chosen to balance the trade-off between spatial resolution and robustness against noise.

##### 1.2.2 Sub-Pixel Estimation and Correlation

For sub-pixel accuracy in the displacement measurement, we employed the Gauss 2x3-point estimator. This method fits a Gaussian function to the cross-correlation peak to estimate the displacement vector with sub-pixel precision. The displacement $\mathbf{d}(\mathbf{x})$ can be expressed as:

$$\mathbf{d}(\mathbf{x}) = \frac{\partial \Phi}{\partial \mathbf{x}} \bigg/ \frac{\partial^2 \Phi}{\partial \mathbf{x}^2}, \tag{4}$$

where $\Phi$ represents the cross-correlation function. The peak of this function is iteratively refined to achieve higher precision.

##### 1.2.3 Post-Processing and Validation

To ensure the reliability of the PIV results, post-processing steps were applied. Vector validation routines, including outlier detection and replacement, were performed using a threshold of 8 times the standard deviation. Additionally, a local median threshold of 3 was applied to detect and correct erroneous vectors. This validation step is crucial for eliminating spurious vectors and improving the overall quality of the velocity field.

##### 1.2.4 PIV Velocity Field Calculation

The final velocity field $\mathbf{v}(\mathbf{x})$ is calculated by dividing the measured displacement $\mathbf{d}(\mathbf{x})$ by the time interval $\Delta t$ between frames:

$$\mathbf{v}(\mathbf{x}) = \frac{\mathbf{d}(\mathbf{x})}{\Delta t}. \tag{5}$$

This velocity field provides a detailed representation of the flow dynamics at each interrogation window's center.

#### 1.3 Grid Averaging and Method Comparison

For comparative analysis between PIV and optical flow, both velocity fields need to be averaged over a uniform grid to facilitate direct comparison.

##### 1.3.1 Grid Averaging Process

The grid averaging process involves interpolating the PIV and optical flow velocity data onto a common grid. Given a grid size $G$, the flow field is averaged over each grid cell. The velocity components $u$ and $v$ are computed as:

$$u_{avg}(i, j) = \frac{1}{n} \sum_{k=1}^{n} u_k, \tag{6}$$

$$v_{avg}(i, j) = \frac{1}{n} \sum_{k=1}^{n} v_k, \tag{7}$$

where $u_k$ and $v_k$ are the velocity components within the grid cell $(i, j)$, and $n$ is the number of vectors within the cell. This averaging smooths the velocity field, reducing noise and allowing for a clearer comparison of the flow structures between methods.

##### 1.3.2 Comparative Analysis

To compare the performance of the PIV and optical flow methods, we calculate three key metrics: relative speed errors, relative orientation errors, and cross-correlation between the methods and ground truth.

**Relative Speed Error** The relative speed error quantifies the discrepancy in the magnitudes of the velocity vectors between the method under evaluation and the ground truth. It is defined as:

$$\text{Relative Speed Error} = \frac{|V_{\text{method}} - V_{\text{gt}}|}{V_{\text{gt}}}, \tag{8}$$

where $V_{\text{method}}$ and $V_{\text{gt}}$ are the magnitudes of the velocity vectors from the method under consideration (either PIV or optical flow) and the ground truth, respectively. This metric highlights the magnitude difference, providing insight into how accurately each method estimates the speed of the flow.

**Relative Orientation Error** The relative orientation error assesses the angular deviation between the estimated velocity vector from the method and the ground truth vector. It is calculated as follows:

$$\text{Relative Orientation Error} = \cos^{-1}\left(\frac{\mathbf{v}_{\text{method}} \cdot \mathbf{v}_{\text{gt}}}{|\mathbf{v}_{\text{method}}||\mathbf{v}_{\text{gt}}|}\right), \tag{9}$$

where $\mathbf{v}_{\text{method}}$ and $\mathbf{v}_{\text{gt}}$ are the velocity vectors obtained from the method and the ground truth, respectively. This metric measures the angular difference between the vectors, indicating how well each method captures the direction of the flow.

**Cross-Correlation Analysis** To evaluate the temporal consistency and overall agreement of the methods with the ground truth, we compute the cross-correlation of the mean speed across frames. The cross-correlation between the method's mean speed and the ground truth's mean speed is defined as:

$$\text{Cross-Correlation} = \frac{\sum(V_{\text{method}} - \bar{V}_{\text{method}})(V_{\text{gt}} - \bar{V}_{\text{gt}})}{\sqrt{\sum(V_{\text{method}} - \bar{V}_{\text{method}})^2 \sum(V_{\text{gt}} - \bar{V}_{\text{gt}})^2}}, \tag{10}$$

where $V_{\text{method}}$ and $V_{\text{gt}}$ are the mean speeds of the method and ground truth for a given frame, and $\bar{V}_{\text{method}}$ and $\bar{V}_{\text{gt}}$ are the overall mean speeds across all frames. This metric assesses how well the trends in the method's velocity estimates correlate with those of the ground truth over time.

##### 1.3.3 Benchmarking

We benchmarked our custom optical flow algorithm with openPIV package from python. We used experimental data from the our previous work [1]. The dataset consist of densely labeled Alexa-647 MTs and 1 µm tracer beads were sequentially imaged every 8 seconds using fluorescence filter or brightfield microscopy. We performed PIV on the tracer beads data as the ground truth and performed PIV and image flow on the densely labeled data. We noticed that our image flow algorithm performed better in calculating both the magnitude of the velocity field and the orientation of the velocity vectors as shown in supplemental figure 1.

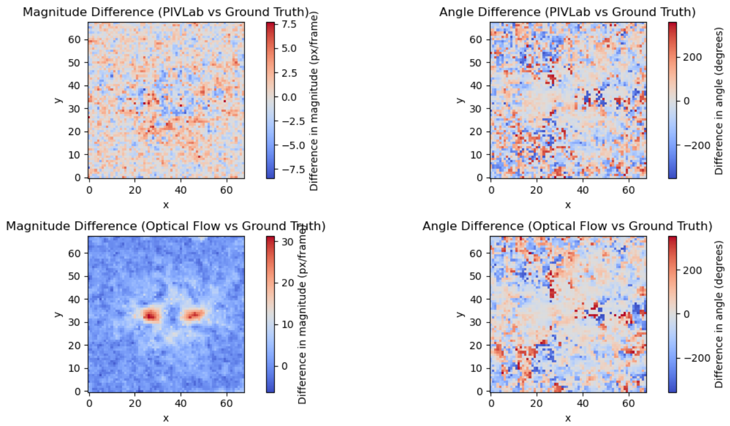


Figure 1: **Comparison of Magnitude and Angle Differences between PIV/Optical Flow and Ground Truth for Frame 20.** The top row shows the **PIVLab** method compared to the ground truth: (left) magnitude difference and (right) angle difference. The bottom row shows the **Optical Flow** method compared to the ground truth: (left) magnitude difference and (right) angle difference. The magnitude difference is measured in pixels per frame, and the angle difference is in degrees. The PIVLab results exhibit less pronounced differences in magnitude and a scattered pattern of angle differences. In contrast, the optical flow method shows localized areas with significant magnitude differences but more uniform angle discrepancies.

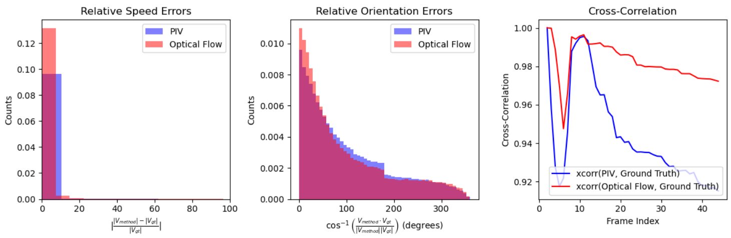


Figure 2: **Histograms of Relative Speed and Orientation Errors, and Cross-Correlation for PIV and Optical Flow Methods.** From left to right: (1) The **relative speed errors** histogram shows the distribution of speed discrepancies normalized by the ground truth velocity. (2) The **relative orientation errors** histogram shows the distribution of angular discrepancies between the estimated and ground truth vectors. (3) The **cross-correlation** plot between the methods' mean speed and the ground truth's mean speed over 45 frames. Higher cross-correlation values indicate better temporal consistency with the ground truth. The optical flow method (red) maintains a high and stable correlation, while the PIV method (blue) shows a decline in correlation over time.

#### 2 Microtubule Concentration Calibration

To establish a quantitative relationship between microtubule concentration and image intensity, we developed a calibration curve using a controlled experimental setup. This calibration allows us to convert observed fluorescence intensities in our main experiments to microtubule densities.

Code implementation in figure_1 _density.ipynb

##### 2.1 Experimental Procedure

We prepared microtubule solutions at eight different concentrations. Each concentration was loaded into a separate flow cell. For each concentration, we captured 10 fluorescence microscopy images at different locations within the flow cell.

##### 2.2 Data Analysis

For each image, we calculated the average pixel intensity. We then computed the mean intensity and standard deviation across the 10 images for each concentration. The known microtubule concentrations were converted to densities (MT/$\mu$m$^2$) based on the flow cell dimensions.

##### 2.3 Calibration Curve

We plotted the average pixel intensity against the microtubule density and performed a linear regression. The resulting calibration equation is:

$$I = 143.05\rho - 39.76 \tag{11}$$

where:

- $I$ is the average pixel intensity
- $\rho$ is the microtubule density in MT/$\mu$m$^2$

#### 3 Velocity Correlation Function and Correlation Length Calculation

To quantify the spatial coherence of the velocity field in our active microtubule network, we computed the two-point velocity correlation function and derived the correlation length. This analysis provides insights into the length scale over which the system exhibits coordinated motion.

Code implementation in figure_1 _correlation.ipynb

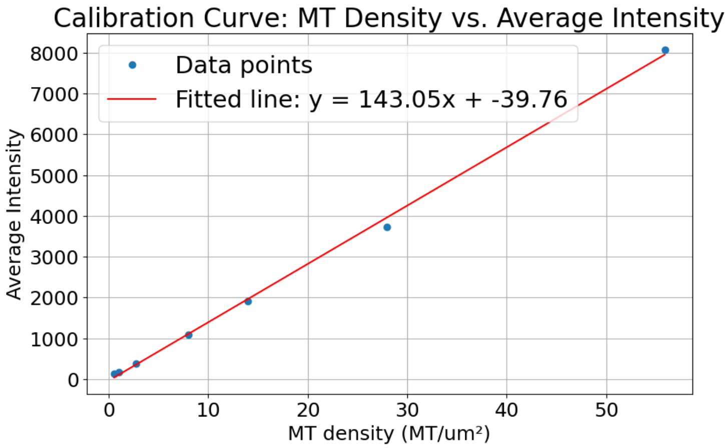


Figure 3: **Microtubule density calibration curve**

##### 3.1 Velocity Correlation Function

The two-point velocity correlation function $C(r)$ is defined as:

$$C(r) = \frac{\langle \mathbf{v}(\mathbf{r}_0) \cdot \mathbf{v}(\mathbf{r}_0 + \mathbf{r}) \rangle}{\langle |\mathbf{v}(\mathbf{r}_0)|^2 \rangle} \tag{12}$$

where $\mathbf{v}(\mathbf{r})$ is the velocity vector at position $\mathbf{r}$, and the angle brackets denote an ensemble average over all positions $\mathbf{r}_0$ and time. Velocity vectors were computed based on Supplemental Information 1.1 In practice, we compute this correlation function as follows:

We select a region of interest in the center of the image, spanning $\pm 100$ pixels in the y-direction. We compute the mean velocity field by averaging over frames 90 to 110. For each frame, we calculate the velocity fluctuations: $\delta \mathbf{v}(\mathbf{r}) = \mathbf{v}(\mathbf{r}) - \langle \mathbf{v}(\mathbf{r}) \rangle$ We then compute the correlation between a reference point (rightmost column of the ROI) and all other points to its left:

$$C(r) = \frac{\langle \delta \mathbf{v}(\mathbf{r}_{\text{ref}}) \cdot \delta \mathbf{v}(\mathbf{r}_{\text{ref}} + \mathbf{r}) \rangle}{\sqrt{\langle |\delta \mathbf{v}(\mathbf{r}_{\text{ref}})|^2 \rangle \langle |\delta \mathbf{v}(\mathbf{r}_{\text{ref}} + \mathbf{r})|^2 \rangle}} \tag{13}$$

We average this correlation over all frames to obtain the final correlation function.

##### 3.2 Correlation Length Calculation

The correlation length $L_c$ is a measure of the distance over which velocities remain correlated. We use two methods to calculate $L_c$, depending on the behavior of the correlation function:

For short incubation times (local phase): We fit the correlation function to an exponential decay:

$$C(r) = Ae^{-r/L_c} \tag{14}$$

where $A$ is the amplitude and $L_c$ is the correlation length. We use non-linear least squares fitting to determine $L_c$.

For long incubation times (global phase): The correlation function exhibits more complex behavior, often becoming negative at larger distances. In this case, we define $L_c$ as the distance at which the correlation function first drops to $1/e$ of its initial value:

$$C(L_c) = \frac{1}{e}C(0) \tag{15}$$

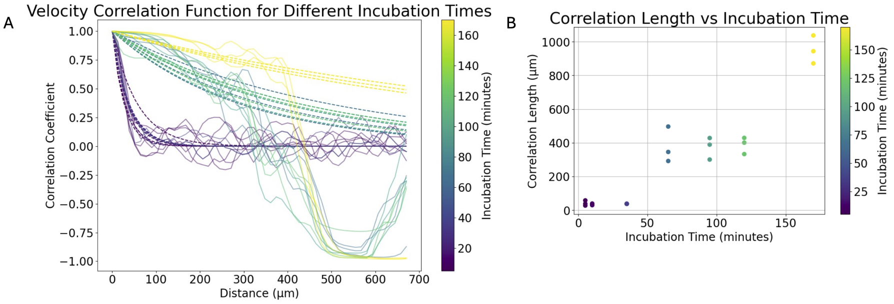


Figure 4: **Exponential decay fitted to the two-point correlation function, and the corresponding correlation length l**

### 4 Flux Analysis

Code implementation in figure_1 _flux.ipynb

#### 4.1 Elliptical Region of Interest (ROI) Definition

We define an elliptical ROI using the equation:

$$\left(\frac{x - x_c}{a}\right)^2 + \left(\frac{y - y_c}{b}\right)^2 \leq 1 \tag{16}$$

where $(x_c, y_c)$ is the center of the ellipse, and $a$ and $b$ are the semi-major and semi-minor axes, respectively.

#### 4.2 Boundary Point Selection

We use a k-d tree algorithm to efficiently find the nearest points on the ROI boundary to a set of evenly distributed points on a perfect ellipse. This ensures uniform sampling around the ROI while adhering to the pixelated nature of the image.

##### 4.3 Flux Calculation

For each boundary point $i$ at time $t$, we calculate the flux $F_i(t)$ as:

$$F_i(t) = \pm v_x(i,t) \cdot \rho(i,t) \cdot \Delta x \tag{17}$$

where $v_x(i,t)$ is the x-component of the velocity, $\rho(i,t)$ is the microtubule density, and $\Delta x$ is the pixel size. The sign is positive for points on the left half of the ellipse and negative for points on the right half.

##### 4.4 Density Calibration

We convert intensity to microtubule density using the linear relationship using the equation obtained in section 2.

##### 4.5 Data Smoothing

We apply Gaussian smoothing to the flux data:

$$F_{\text{smooth}}(x) = \int F(t) \cdot G(x-t)dt \tag{18}$$

where $G(x)$ is a Gaussian function with standard deviation $\sigma = 2$.

##### 4.6 Cumulative and Total Flux

Cumulative flux for each point $i$: $C_i = \sum_t F_i(t)$ Total flux at time $t$: $T(t) = \sum_i F_i(t)$

##### 4.7 Velocity Field Interpolation

We interpolate the velocity field to the ROI boundary points using a grid-based approach:

$$v(x,y) \approx v(\lfloor x/g \rfloor \cdot g, \lfloor y/g \rfloor \cdot g) \tag{19}$$

where $g$ is the grid size (30 pixels in this case).

#### 5 Fluorescence Recovery After Photobleaching (FRAP)

Fluorescence Recovery After Photobleaching (FRAP) is a technique used to study the dynamics of molecular diffusion and interactions within cellular environments. In a FRAP experiment, a region of interest (ROI) within a fluorescently labeled sample is photobleached using a high-intensity laser, and the subsequent recovery of fluorescence within the bleached area is monitored over time. The rate of fluorescence recovery provides insights into the mobility and

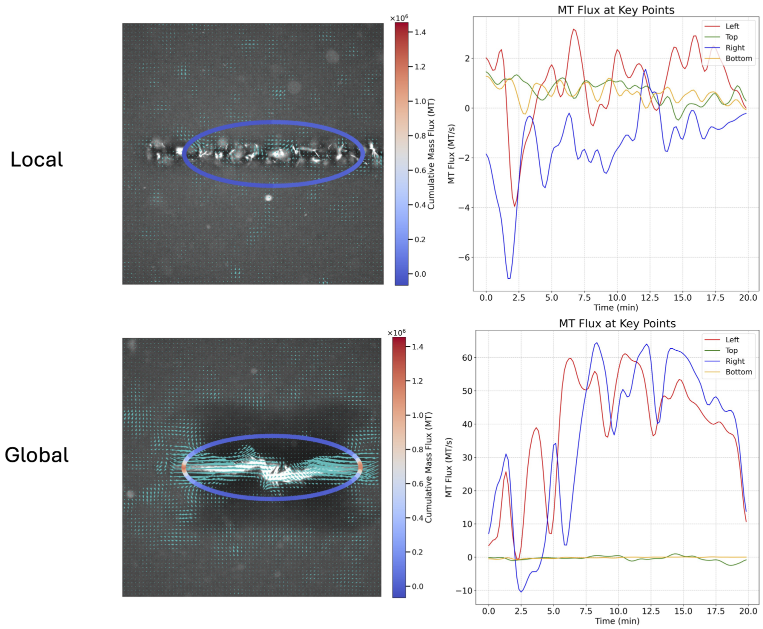


Figure 5: **Microtubule flux at vertices and co-vertices of the ROI in representative global and local phase.**

binding characteristics of the molecules in the ROI. Since we cannot directly image and measure the microtubule structures in the experiments, we employed FRAP to measure for the microtubule structure size. Code implementation in figure_2 _FRAP.ipynb

#### 5.1 Data Normalization

Normalization of the fluorescence intensity values is performed to account for variability in initial fluorescence levels and photobleaching efficiency. The normalization is defined as:

$$I_{\text{normalized}}(t) = \frac{I(t) - I_{\text{min}}}{I_{\text{max}} - I_{\text{min}}}, \tag{20}$$

where $I(t)$ is the intensity at time $t$, $I_{\text{min}}$ is the minimum intensity, and $I_{\text{max}}$ is the maximum intensity observed in the recovery phase. This normalization ensures that the fluorescence intensity values range between 0 and 1.

#### 5.2 Half-time Recovery Calculation

The half-time recovery ($\tau_{1/2}$) is a critical parameter in FRAP analysis, representing the time required for the fluorescence intensity to recover to half of its

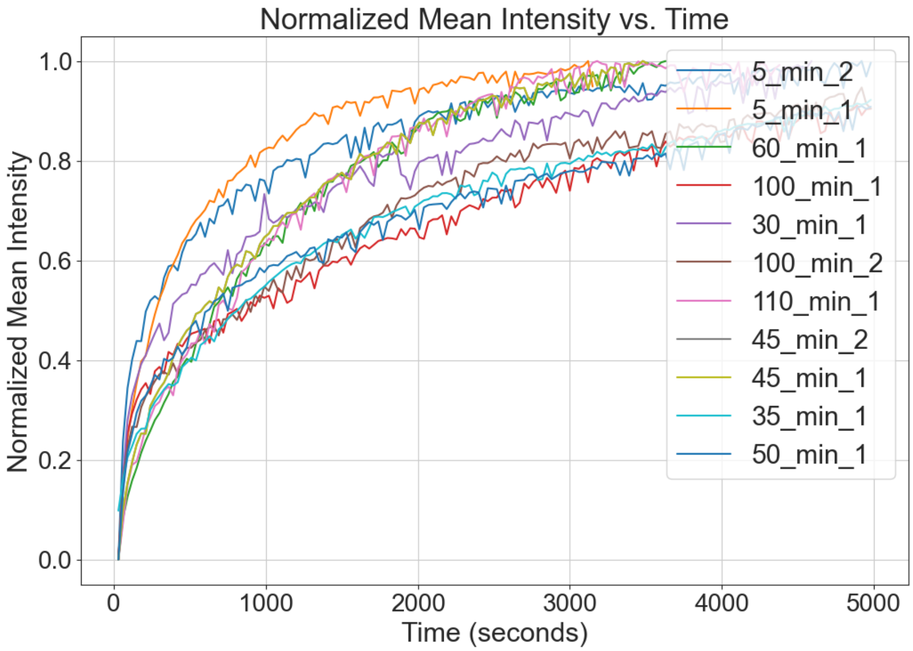


Figure 6: **Fluorescence Intensity During Recovery**

final value. Mathematically, it is determined as:

$$\tau_{1/2} = \min\left\{t \mid I_{\text{normalized}}(t) \geq 0.5\right\}. \tag{21}$$

This value is identified by finding the time point at which the normalized intensity reaches 50

#### 5.3 Diffusion Coefficient Calculation

The diffusion coefficient $D$ quantifies the rate at which molecules diffuse within the bleached area. It is derived from the half-time recovery using the following equation from Axelrod's paper:

$$D = \frac{w^2}{4\tau_{1/2}}\gamma_D, \tag{22}$$

where $w$ is the radius of the laser beam used for photobleaching, and $\gamma_D = 0.88$ is a geometric correction factor for circular beams. The radius $w$ is calculated as:

$$w = \text{resolution} \times \text{pixels}, \tag{23}$$

with the resolution of the optical setup being $0.294\,\mu\text{m/pixel}$ and the number of pixels being 390, giving:

$$w = 0.294\,\mu\text{m/pixel} \times 390\,\text{pixels} = 114.66\,\mu\text{m}. \tag{24}$$

Thus, the diffusion coefficient is:

$$D = \frac{(114.66\,\mu\text{m})^2}{4\tau_{1/2}} \times 0.88. \tag{25}$$

### 6 Stokes Force Calculation

To calculate the forces exerted by the fluid on tracer particles, we employed Stokes' law. This method involved tracking the motion of tracer beads within the fluid, determining their velocities, and then calculating the forces based on the fluid's viscosity and the particles' radii.

#### 6.1 Particle Tracking

We utilized the same particle tracking code as described in the previous section, but with 10 µm tracer beads introduced into the system. The positions of these beads, $\mathbf{x}_i(t)$, were tracked over time $t$, and their velocities, $\mathbf{v}_i(t)$, were computed by differentiating their trajectories with respect to time:

$$\mathbf{v}_i(t) = \frac{d\mathbf{x}_i(t)}{dt}. \tag{26}$$

#### 6.2 Velocity Calculation

The velocities of the tracer beads were initially calculated in micrometers per second ($\mu$m/s). These velocities were then converted to meters per second (m/s) for use in the Stokes' law calculation:

$$v_{i,\text{m/s}} = v_{i,\mu\text{m/s}} \times 10^{-6}. \tag{27}$$

#### 6.3 Stokes' Law

Stokes' law relates the force $F$ exerted by a viscous fluid on a spherical particle moving through the fluid to the particle's radius $r$, the fluid's viscosity $\eta$, and the particle's velocity $v$. The equation is given by:

$$F = 6\pi\eta r v. \tag{28}$$

For our system, we used the following parameters:

- Radius of the tracer particles: $r = 6.5 \times 10^{-6}$ meters.
- Viscosity of the fluid: $\eta = 2$ Pa.s.

The calculated forces for each particle were then averaged to obtain a representative value of the force in the system. The results, including velocities in both $\mu$m/s and m/s, and the corresponding forces in Newtons (N), were compiled into a DataFrame for further analysis and plotting.

##### 6.4 Example Calculation

For clarity, an example calculation is provided below. Given a tracer bead velocity of $10\,\mu$m/s, the force exerted on the particle is calculated as follows:

$$\begin{aligned} v_{i,\mathrm{m/s}} &= 10 \times 10^{-6}\,\mathrm{m/s}, \\ F_i &= 6\pi \times 2 \times 6.5 \times 10^{-6} \times 10 \times 10^{-6}, \\ F_i &= 2.44 \times 10^{-9}\,\mathrm{N}. \end{aligned}$$

### 7 Percolation Analysis

##### 7.1 Mapping active-gel connectivity onto ER percolation

In the main text we coarse-grain the microtubule–kinesin network into a *length-controlled* random graph: each microtubule bundle is a node; an *undirected* edge is drawn whenever any two bundles overlap or cross-link within a chemically short range. Code implementation of the simulation in "percolation.ipynb" an simpler version could be found If we assume (i) bundles are well mixed in 2-D chambers and (ii) the probability of a contact scales only with their mean length $\langle L\rangle$, then the adjacency matrix becomes *dense* and *uniform.* Under this assumption the ensemble is analytically equivalent to the classical Erdős–Rényi (ER) model

$$\mathcal{G}(n,q), \quad q = \frac{c}{n}, \qquad c = 1 + \delta,$$

where $n$ is the number of bundles, $q$ the edge probability and $c$ the mean degree. The control parameter is the reduced distance to criticality $\delta = c - 1$.

##### 7.2 Order parameter and susceptibility

For every generated graph we label connected components with `scipy.sparse.csgraph.connected_component` Let $N_{\mathrm{GC}}$ be the size of the largest cluster (the *giant component*). We monitor two standard observables:

$$S = N_{\mathrm{GC}}/n, \tag{29}$$

$$\chi = \frac{1}{n} \sum_{\alpha \neq \mathrm{GC}} N_\alpha^2, \tag{30}$$

where the sum runs over all clusters except the giant one. $S$ plays the rôle of a magnetisation-like order parameter, while $\chi$ is the cluster-size susceptibility.

##### 7.3 Finite-size scaling protocol

**System sizes.** We simulate three graph sizes,

$$n \in \{6\,400,\ 12\,800,\ 25\,600\},$$

chosen so that $n^{1/3}$ is an integer power of two, simplifying the asymptotic window selection.

**Asymptotic window.** Mean-field theory predicts $\delta_{\min} \sim n^{-1/3}$ and we restrict all power-law fits to

$$|\delta| \in \left[n^{-1/3},\ 0.04\right].$$

The upper bound 0.04 is well inside the critical regime yet far from finite-size saturation.

**Replicas and averaging.** For each $(n, \delta)$ pair we generate $N_{\text{rep}} = 60$ independent graphs. Sample means $\overline{S}$ and $\overline{\chi}$ are then used in the fits.

**Exponent extraction.** In the ER universality class

$$S \sim \delta^{\beta}, \qquad \chi \sim |\delta|^{-\gamma}, \qquad \beta = \gamma = 1,$$

for $\delta > 0$ and $\delta < 0$ respectively. We estimate slopes with a simple log–log linear regression; zeros and infinities are masked.

**Data collapse.** Setting $\nu = \frac{1}{2}$ (exact mean-field value) we rescale

$$x = \delta\, n^{1/3}, \quad y = S\, n^{1/3},$$

and verify that all sizes fall on a single master curve (Fig. S1).

**Numerical result.** Table 1 summarises the fitted exponents. The statistical uncertainty (standard error from the linear fit) is $\lesssim$0.05 for all $n$.

Table 1: Numerical exponents for $S$ and $\chi$.

| $n$ | $\beta$ | $\gamma$ |
|---|---|---|
| 6 400 | 1.04 | 0.96 |
| 12 800 | 1.01 | 1.02 |
| 25 600 | 0.98 | 1.01 |

#### 7.4 Gamma-length phase diagram

To emulate the experimentally observed bundle-length distribution we draw filament lengths $\ell$ from a *clamped* Gamma law,

$$\ell \;\sim\; \Gamma(k, \theta)\big[\,0, \ell_{\max} = 25\,\mu\text{m}\big],$$

parameterised by the shape $k$ and scale $\theta$ of the underlying (unclamped) Gamma density. For every $(k, \theta)$ pair we compute the *tail probability*

$$q(k, \theta) \;=\; 1 - \,F_{\Gamma}\big(\ell_{\text{thr}} = 9\,\mu\text{m};\ k, \theta\big),$$

where $F_\Gamma$ is the cumulative distribution function of the *unclamped* Gamma. The threshold $\ell_{\rm thr}{=}9\,\mu$m corresponds to the minimum overlap length at which two bundles are experimentally observed to cross-link with high probability.

We then feed the edge probability $q$ directly into an Erdős–Rényi graph $\mathcal{G}(n,q)$ with $n = 25\,600$ Instead of Monte-Carlo sampling we exploit the $n \to \infty$ closed-form solution

$$S \;=\; 1 + \frac{W\bigl(-c\,e^{-c}\bigr)}{c}, \qquad c = n\,q,$$

where $W$ is the Lambert $W$ function. This expression is exact for ER graphs and makes the $120 \times 120$ parameter scan complete in under one second on a laptop.

Figure 2I of the main text plots the resulting surface $S(k,\theta)$ as a heat-map. The color jump delineates a thin percolation front: increasing either the shape or the scale parameter by $\Delta k \lesssim 0.2$ or $\Delta\theta \lesssim 0.2$ is enough to nucleate a system-spanning (giant) component.

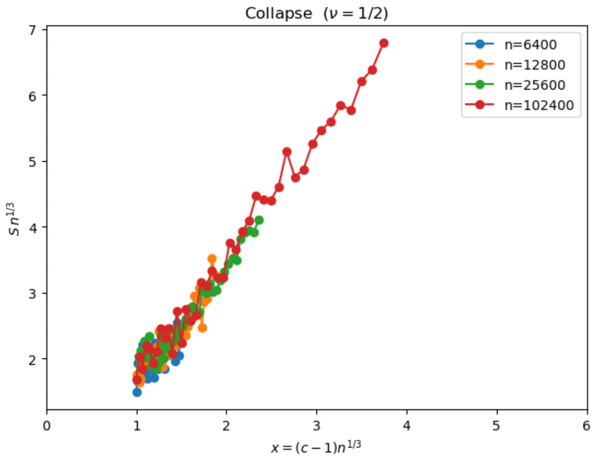


Figure 7: **Finite-size scaling collapse of the giant-component order parameter.** Symbols denote Erdős–Rényi graphs with $n$ = 6,400 (blue), 12,800 (orange), 25,600 (green) and 102,400 (red). Only data inside the asymptotic window $n^{-1/3} \le |\delta| \le 0.08$ (see Supplementary Note S3) are plotted, so the rescaled axis begins at $x_{\min} \approx 1$. After the mean-field rescaling $x = (c-1)\,n^{1/3}$ and $y = S\,n^{1/3}$ all sizes fall on a single master curve, confirming the expected exponents $\nu = \frac{1}{2}$ and $\beta = 1$.

#### 7.5 Reproducibility notes

- All simulations were run with `Python 3.10`, `SciPy 1.11`, `NumPy 1.26`, `scikit-learn 1.4`, and `Matplotlib 3.8`.

- A fully annotated Jupyter notebook (`percolation.ipynb`) accompanies this file. A lighter Colab version is linked in the main text.
- Random streams are seeded with `np.random.default_rng(seed=2024)` for bit-wise reproducibility.
- Runtime on a laptop (Apple M1, 8 cores) is $\sim 30$s for the scaling analysis and $\sim 20$s for the phase diagram.

### 8 Cytosim Simulation

#### 8.1 Mathematical Background and Cytosim Framework

Cytosim is a powerful software package designed to simulate the collective dynamics of cytoskeletal fibers, such as microtubules and actin filaments [2]. It uses a coarse-grained representation of fibers and includes various biophysical processes, such as fiber flexibility, polymerization, depolymerization, and motor protein interactions. The mathematical framework underlying Cytosim is based on Langevin dynamics, which describes the motion of fibers and motors in a viscous medium subject to thermal fluctuations. In Cytosim, fibers are modeled as inextensible elastic rods, which are discretized into segments. The dynamics of each segment $i$ is governed by the Langevin equation:

$$\gamma \frac{d\mathbf{r}_i}{dt} = \mathbf{F}_i^{\text{elastic}} + \mathbf{F}_i^{\text{motor}} + \mathbf{F}_i^{\text{thermal}} + \mathbf{F}_i^{\text{constraint}} \tag{31}$$

where $\gamma$ is the drag coefficient, $\mathbf{r}_i$ is the position of segment $i$, and $\mathbf{F}_i$ represents the forces acting on the segment. The elastic forces $\mathbf{F}_i^{\text{elastic}}$ are derived from the bending and twisting energies of the fibers, while the motor forces $\mathbf{F}_i^{\text{motor}}$ are described by a stochastic binding and unbinding kinetics. The thermal forces $\mathbf{F}_i^{\text{thermal}}$ are modeled as Gaussian white noise, and the constraint forces $\mathbf{F}_i^{\text{constraint}}$ ensure the inextensibility of the fibers and the connectivity between segments. The Langevin equation is solved numerically using a semi-implicit integration scheme:

$$\mathbf{r}_i(t + \Delta t) = \mathbf{r}_i(t) + \frac{\Delta t}{\gamma} \left( \mathbf{F}_i^{\text{elastic}} + \mathbf{F}_i^{\text{motor}} + \mathbf{F}_i^{\text{thermal}} + \mathbf{F}_i^{\text{constraint}} \right) \tag{32}$$

where $\Delta t$ is the time step. This scheme ensures numerical stability and allows for the efficient simulation of large systems with many fibers and motors.

#### 8.2 Generating Cytosim Configuration Files

To run simulations in Cytosim, users need to provide configuration files that specify the properties and initial conditions of the cytoskeletal components. In this study, we focus on the role of microtubule length distribution in the collective dynamics of microtubule-kinesin systems. We generate Cytosim configuration files with different microtubule length distributions based on the gamma distribution, which is characterized by two parameters: shape ($k$) and scale ($\theta$).

We define a function `generate_microtubule_lengths(shape, scale, total_length)` that generates a set of microtubule lengths following a gamma distribution with the specified shape and scale parameters. The function also takes a `total_length` argument, which determines the target sum of all generated microtubule lengths. The generated lengths are rounded to one decimal place and constrained to be between 1 and 25 units, which is a biologically relevant range for microtubule lengths.

The generated microtubule lengths are then used to create microtubule objects in the Cytosim configuration files. For each unique length value in the generated lengths list, a microtubule object is created with the corresponding length and the number of microtubules with that length. For example:

```
new 5 microtubule
{
length = 1.5
}
new 3 microtubule
{
length = 2.3
}
```

In this example, there are 5 microtubules with a length of 1.5 and 3 microtubules with a length of 2.3.

To explore the effect of the gamma distribution parameters on the microtubule lengths and the resulting collective dynamics, we generate config files for two different scenarios: a 1D parameter sweep and a 2D parameter sweep. In the 1D sweep, we fix the shape parameter and vary the scale parameter, while in the 2D sweep, we vary both parameters simultaneously. Code implementation can be found in 1d-phase-diagram.ipynb and 2d-phase-diagram.ipynb, and manual-config.ipynb

### 9 Microtubule Aster Tracking

Tracking microtubule asters involves detecting and tracking their positions over time in image sequences. To improve detection accuracy, variable thresholding is applied, which adjusts the thresholding percentile based on the image characteristics, ensuring consistent detection across frames with varying intensity distributions. "figure_4_aster _traj.ipynb"

#### 9.1 Preprocessing with Variable Thresholding

Variable thresholding adapts the threshold value dynamically by calculating the percentile of pixel intensities in each frame. This method helps in distinguishing microtubules from the background more effectively. The preprocessing steps are:

1. Calculate the threshold value as a specified percentile of pixel intensities.

2. Create a binary mask where pixels above the threshold are set to 255 (white) and others to 0 (black).
3. Apply binary dilation to enhance the visibility of microtubule structures.

For different segments of the image sequence, different percentiles are used:

- The first 20 frames use a 99.4 percentile threshold.
- Frames 21 to 30 use a 99.5 percentile threshold.
- Frames 31 and beyond use a 99.97 percentile threshold.

##### 9.2 Aster Tracking

After preprocessing, the TrackPy library is used to detect and track microtubule asters. The tracking steps are:

1. Detect particles in each frame using a bandpass filter.
2. Link particles across frames to form trajectories based on proximity and predicted motion.
3. Calculate velocities by differentiating trajectories with respect to time.

This method allows for accurate tracking of microtubule asters, providing valuable data on their dynamics under different experimental conditions.

##### 9.3 Aster Area Calculating

Aster area can be computed by calculating the area above the intensity threshold. Implemented in "figure_4_aster_size.ipynb"

#### 10 Cell and Bead Tracking and Velocity Calculation

We tracked cells and beads in brightfield images to analyze their movements and calculate associated physical quantities such as velocity, force, work, and power. The tracking was performed using a similar approach to particle tracking. Code implementation in "figure_4_taj.ipynb"

##### 10.1 Cargo Detection and Tracking

Cargo were imaged using brightfield microscopy. The detection and tracking process involved the following steps:

1. **Detection:** Cargo was identified in each frame based on intensity and morphological criteria.

2. **Linking:** Detected cargo was linked across frames to form trajectories, ensuring that each cell/bead's movement was accurately captured over time.

The position $\mathbf{x}_i(t)$ of cargo $i$ at time $t$ was recorded, allowing for the calculation of displacement and velocity.

#### 10.2 Velocity Calculation

The velocity $\mathbf{v}_i(t)$ of cargo $i$ was calculated by differentiating its trajectory with respect to time.

$$\text{velocity} = \text{velocity} \times 0.43\,\mu\text{m/pixel} \tag{33}$$

#### 10.3 Tracer Bead Force Calculation

The force exerted by or on the cargo was calculated using Stokes' law, previously described in section 5. In short, we calculate the force as the force experienced by a spherical particle moving through a viscous fluid to its velocity:

$$F = 6\pi\eta r v, \tag{34}$$

where:

- $F$ is the force,
- $\eta$ is the viscosity of the fluid,
- $r$ is the radius of the cell,
- $v$ is the velocity of the cell.

In this case, we used a radius $r = 6.5 \times 10^{-6}$ meters (6.5 µm) and fluid viscosity $\eta = 2 \times 10^{-3}$ Pa.s. The velocity $v$ was converted from µm/s to m/s using the pixel-to-meter conversion factor.

#### 10.4 Work and Power Calculation

The work done by the moving the cargo was calculated by integrating the force over the displacement for each time step:

$$\text{work\_done} = F \cdot \text{displacement}, \tag{35}$$

where the displacement was converted from pixels to meters.

The total work done over the entire observation period was obtained by summing the work done at each time step.

$$\text{total\_work\_done} = \sum \text{work\_done} \tag{36}$$

The power, which is the rate of doing work, was calculated by dividing the work done at each time step by the corresponding time interval:

$$\text{power} = \frac{\text{work\_done}}{\Delta t}, \tag{37}$$

where $\Delta t$ is the time between frames (1/8 seconds).

The total power at each time step was then summed to obtain the overall power profile as a function of time, which was plotted to visualize the energy expenditure of the cells.

$$\text{total\_power} = \sum \text{power} \tag{38}$$

The power was plotted against time to show how the energy expenditure varied over the course of the experiment.

### 11 Experimental Setup

#### 11.1 Light-induced kinesin expression and purification

We constructed two chimeras of D. melanogaster kinesin K401: K401-iLID and K401-micro as previously described (Addgene 122484 and 122485) [3]. In short, for the K401-iLiD plasmid we inserted iLID with a His tag after the C-terminus of K401. For the K401-micro plasmid, we inserted K401 between the His-MBP and the micro. The MBP domain is needed to ensure the microdomain remains fully functional during expression [4]. After the expression, the MBP domain can be cleaved of by TEV protease.

For protein expression, we transformed the plasmids to BL21(DE3)pLysS cells. The cells were grown in LB and induced at with 1mM IPTG at 18°C for 16 hours after reaching OD 0.6. The cells were then pelleted at 4000G and resuspended in lysis buffer (50 mM sodium phosphate, 4 mM MgCl2, 250 mM NaCl, 25 mM imidazole, 0.05mM MgATP, 5 mM BME, 1 mg/ml lysozyme and 1 tablet/50 mL of Complete Protease Inhibitor). After a 1-hour incubation with stirring, the lysate was passed through a 30kPSI cell disruptor. The lysate was clarified at 30,000 G for 1 hour. The supernatant was then incubated with Ni-NTA agarose resin for 1 hour. The lysate/Ni-NTA mixture was loaded into a chromatography column and washed three times with wash buffer (Lysis buffer with no lysozyme nor complete EDTA tablet), and eluted with 500mM imidazole. Protein elutions were dialyzed overnight using 30 kDa MWCO membrane against 50 mM sodium phosphase, 4 mM MgCl2, 250 mM NaCl, 0.05 mM MgATP, and 1 mM BME. For the K401-micro elution, we added TEV protease at a 1:25 mass ratio to remove the MBP domain. Then, we used centrifugal filters to exchange to pH 6.7 protein storage buffer (50 mM imidazole, HCl for pH balancing, 4 mM MgCl2, 2 mM DTT, 50 µM MgATP, and 36% sucrose). Proteins were then aliquoted and flash frozen in LN2 and stored under -80°C.

#### 11.2 Microtubule Polymerization and Length Distribution

Fluorescent microtubule polymerization was previously described [3]. In short, we used a protocol based on one found on the Dogic lab homepage. The procedure began by preloading and starting a 37℃ water bath. GMP-cpp, reagents, and tubes were cooled on ice. A 20 mM DTT solution was prepared using Pierce no-weigh format, and a GMP mixture consisting of M2B, DTT, and GMP-cpp was made and stored on ice. The ultracentrifuge and rotor were pre-cooled to 4℃. Tubulin (20 mg/mL) and labeled tubulin (20 mg/mL) were thawed in the water bath until mostly thawed and then cooled on ice. In a cold room, labeled tubulin was added to the stock vial of unlabeled tubulin and mixed gently. The GMP mixture was then added to the tubulin mixture and stirred gently. This combined mixture was pipetted into ultracentrifuge tubes and incubated on ice for 5 minutes before being centrifuged at 90,000 rpm, 4℃ for 8 minutes. The supernatant was carefully collected without disturbing the pellet and transferred to an Eppendorf tube, mixed, and stored on ice. The mixture was incubated in a 37℃ water bath for 1 hour, protected from light. Aliquots were then dispensed into PCR strip tubes, which were spun to collect the fluid at the bottom. Finally, the PCR strips were flash frozen in liquid nitrogen and stored in a -80℃ freezer.

To measure the length distribution of microtubules, we imaged fluorescently labeled microtubules immobilized onto the cover glass surface of a flow cell. The cover glass was treated with a 0.01% solution of poly-L-lysine (Sigma P4707) to promote microtubule binding. The lengths of microtubules were determined by image segmentation. Each microtubule image was normalized and underwent local and global thresholding to correct for non-uniform backgrounds and obtain thresholded images of putative microtubules. Morphological operations were applied to reconnect small breaks in filaments. Objects near the image boundary were removed, and small or circular objects were filtered out based on size and eccentricity thresholds. Potential microtubule crossovers were identified and eliminated by analyzing the angles of lines within the image.

#### 11.3 Sample Chambers for Experiments

Glass slides and coverslips were first cleaned using a series of washes. Slides and coverslips were placed in respective containers, and 2% Hellmanex solution was prepared by mixing 6 mL Hellmanex with 300 mL DI water, heated, and poured into the containers. The containers were sonicated for 10 minutes, followed by three DI water rinses and an ethanol rinse. Ethanol was added to the containers and sonicated again for 10 minutes, followed by another ethanol rinse and three DI water rinses. Next, 0.1 M KOH was added to the containers, sonicated for 10 minutes, and rinsed three times with DI water. The slides were then etched overnight with 5% HCl and rinsed three times with DI water. Clean slides and coverslips were stored in DI water.

For silane coupling, a 2% acrylamide solution was prepared using 40% acrylamide stock solution, and degassed under vacuum. In a chemical hood, 98.5%

ethanol, 1% acetic acid, and 0.5% silane agent were mixed to prepare the silane-coupling solution, which was immediately poured into the containers with the slides and coverslips and incubated at room temperature for 20-30 minutes. The slides were then rinsed once with ethanol, three times with DI water, and baked at 110°C for 30 minutes or 50°C overnight.

For acrylamide polymerization, the degassed 300 mL of 2% acrylamide solution was moved to a stir plate, and 105 µL TEMED and 210 mg ammonium persulfate were added. The solution was immediately poured over the silane-coupled slides and coverslips and left to polymerize overnight at 4°C. Before use, the slides and coverslips were rinsed with DI water and air-dried.

#### 11.4 Reaction Mixture

For the self-organization experiments, K401-micro, K401-iLID, and microtubules were combined into a reaction mixture to achieve final concentrations of approximately 0.1 µM for each motor type and 1.5-2.5 µM for tubulin, referring to protein monomers for K401-micro and K401-iLID constructs, and protein dimers for tubulin. The sample preparation was conducted under dark-room conditions to minimize unintended light activation, using room light filtered to block wavelengths below 580 nm (Kodak Wratten Filter No. 25). The base reaction mixture included a buffer, MgATP as an energy source, glycerol as a crowding agent, pluronic F-127 for surface passivation, and components for oxygen scavenging (pyranose oxidase, glucose, catalase, Trolox, DTT), along with ATP-recycling reagents (pyruvate kinase/lactic dehydrogenase, phosphoenolpyruvic acid). The reaction mixture consisted of 59.2 mM K-PIPES pH 6.8, 4.7 mM MgCl2, 3.2 mM potassium chloride, 2.6 mM potassium phosphate, 0.74 mM EGTA, 1.4 mM MgATP, 10% glycerol, 0.50 mg/mL pluronic F-127, 2.9mg/mL pyranose oxidase, 3.2 mg/mL glucose, 0.086 mg/mL catalase, 5.4 mM DTT, 2.0 mM Trolox, 0.026 units/µL pyruvate kinase/lactic dehydrogenase, and 26.6 mM phosphoenolpyruvic acid.

We note that the sample is sensitive to the buffer pH and mixture incubation time. For our experimental conditions, the mixture pH is around 6.4 and we perform the experiments within 2 hours of constructing the mixture.

#### 11.5 Microscope Setup

We conducted the experiments using an automated widefield epifluorescence microscope (Nikon Ti-2), custom-modified for two additional imaging modes: epi-illuminated pattern projection and LED-gated transmitted light. Light patterns from a programmable DLP chip (EKB Technologies DLP LightCrafter™ E4500 MKII™ Fiber Couple) were projected onto the sample via a user-modified epi-illumination attachment (Nikon T-FL). The DLP chip was illuminated by a fiber-coupled 470 nm LED (ThorLabs M470L3). The epi-illumination attachment featured two light-path entry ports: one for the projected pattern light path and the other for a standard widefield epi-fluorescence light path. These light paths were combined using a dichroic mirror (Semrock BLP01-488R-25).

The magnification of the epi-illumination system was calibrated to ensure the camera's imaging sensor (FliR BFLY-U3-23S6M-C) was fully illuminated when the entire DLP chip was activated. Micro-Manager software, running custom scripts, controlled the pattern projection and stage movement.

#### 11.6 Tracer Particle Method for Measuring Fluid Velocity

To measure the fluid velocity, we employed 1 µm polystyrene beads (Polysciences 07310-15) as tracer particles. The hydrophobic surface of these beads was passivated by incubating them overnight in M2B buffer with 50 mg/ml of pluronic F-127. This incubation process helps to coat the beads with pluronic, which prevents nonspecific interactions with the surrounding fluid and ensures accurate velocity measurements.

Before each experiment, the pluronic-coated beads were thoroughly washed. This was achieved by pelleting the beads through centrifugation and subsequently resuspending them in M2B buffer containing 0.5 mg/ml pluronic. This concentration of pluronic was chosen to match that of the reaction mixture, thereby maintaining consistency in the experimental conditions.

The beads, once prepared, were introduced into the fluid of interest. Their motion was tracked using high-resolution imaging techniques, and their velocities were calculated by analyzing their trajectories over time. This method provides precise measurements of the fluid velocity by leveraging the well-characterized motion of the tracer particles.

#### 11.7 Culturing Jurkat Cells Using RPMI Medium

Jurkat cells, a type of immortalized T lymphocyte cell line, were cultured using RPMI 1640 medium. The RPMI 1640 medium provides the necessary nutrients and environment for the optimal growth and maintenance of Jurkat cells.

The cells were maintained in RPMI 1640 medium supplemented with the following components to enhance cell viability and proliferation:

- 10% fetal bovine serum (FBS) to provide essential growth factors, hormones, and proteins.
- 1% penicillin-streptomycin to prevent bacterial contamination.
- 2 mM L-glutamine to support protein and nucleotide synthesis.

The culturing process involved the following steps:

1. **Cell Seeding:** Jurkat cells were seeded into culture flasks at an initial density of $0.2 \times 10^6$ cells/ml.
2. **Incubation:** The flasks were incubated at 37°C in a humidified atmosphere containing 5% $CO_2$. This environment mimics the physiological conditions of the human body, promoting optimal cell growth.

3. **Medium Change:** The culture medium was replaced every 2-3 days to remove metabolic waste products and replenish nutrients. This was done by gently pelleting the cells through centrifugation, discarding the old medium, and resuspending the cells in fresh RPMI 1640 medium with supplements.

4. **Cell Passage:** When the cells reached a density of $1 \times 10^6$ cells/ml, they were subcultured to maintain optimal growth conditions. This involved diluting the cell suspension with fresh medium and seeding it into new culture flasks.

We also tested cell viability after prolonged incubation in active matter mix using Trypan Blue. Jurkat cells showed minimal cell death during the first 180 minutes of incubation in active matter mixture.

##### 11.8 Active Matter in Oil Emulsion

To prepare the active matter in oil emulsion using the EURx MICELLULA DNA Emulsion kit, follow these steps:

1. **Prepare the Oil Emulsion Mix**

| **Component** | **Volume (µL)** |
|---|---|
| Component 1 | 22 |
| Component 2 | 2 |
| Component 3 | 6 |

2. **Combine the Reaction Mix with the Oil Emulsion Mix**
   - Add 5 µL of the reaction mix to 30 µL of the prepared oil emulsion mix.

3. **Create a Stable Emulsion**
   - Run the tube containing the mixture against a tube rack to create a stable emulsion.
   - Vortex the mixture if necessary to ensure proper emulsification.

#### 12 Code Avilibility

Code can be found at https://github.com/shichenliu97/active-matters-programmed-phase-transition

#### References


[1] Fan Yang, Shichen Liu, Heun Jin Lee, Rob Phillips, and Matt Thomson. Dynamic flow control through active matter programming language. August 2022.

[2] Francois Nedelec and Dietrich Foethke. Collective langevin dynamics of flexible cytoskeletal fibers. *New J. Phys.*, 9(11):427–427, November 2007.

[3] Tyler D Ross, Heun Jin Lee, Zijie Qu, Rachel A Banks, Rob Phillips, and Matt Thomson. Controlling organization and forces in active matter through optically defined boundaries. *Nature*, 572(7768):224–229, August 2019.

[4] G. Guntas, R. A. Hallett, S. P. Zimmerman, T. Williams, H. Yumerefendi, J. E. Bear, and B. Kuhlman. Engineering an improved light-induced dimer (iLID) for controlling the localization and activity of signaling proteins. *Proc. Natl. Acad. Sci. U. S. A.*, 112(1):112–117, 2015.
